# The hepatocyte insulin receptor is required to program rhythmic gene expression and the liver clock

**DOI:** 10.1101/2021.02.05.430014

**Authors:** Tiffany Fougeray, Arnaud Polizzi, Marion Régnier, Anne Fougerat, Sandrine Ellero-Simatos, Yannick Lippi, Sarra Smati, Frédéric Lasserre, Blandine Tramunt, Marine Huillet, Léonie Dopavogui, Lorraine Smith, Claire Naylies, Caroline Sommer, Alexandre Benani, Joel T. Haas, Walter Wahli, Hélène Duez, Pierre Gourdy, Laurence Gamet-Payrastre, Anne-Françoise Burnol, Nicolas Loiseau, Catherine Postic, Alexandra Montagner, Hervé Guillou

## Abstract

In mammalian cells, gene expression is rhythmic and sensitive to various environmental and physiological stimuli. A circadian clock system helps to anticipate and synchronize gene expression with daily stimuli including cyclic light and food intake, which control the central and peripheral clock programs, respectively. Food intake also regulates insulin secretion. How much insulin contributes to the effect of feeding on the entrainment of the clock and rhythmic gene expression remains to be investigated.

An important component of insulin action is mediated by changes in insulin receptor (IR)-dependent gene expression. In the liver, insulin at high levels controls the transcription of hundreds of genes involved in glucose homeostasis to promote energy storage while repressing the expression of gluconeogenic genes. In type 2 diabetes mellitus (T2DM), selective hepatic insulin resistance impairs the inhibition of hepatic glucose production while promoting lipid synthesis. This pathogenic process promoting hyperlipidemia as well as non-alcoholic fatty liver diseases.

While several lines of evidence link such metabolic diseases to defective control of circadian homeostasis, the hypothesis that IR directly synchronizes the clock has not been studied *in vivo*. Here, we used conditional hepatocyte-restricted gene deletion to evaluate the role of IR in the regulation and oscillation of gene expression as well as in the programming of the circadian clock in adult mouse liver.

## INTRODUCTION

The mammalian clock system allows cells to anticipate environmental stimuli and synchronize daily physiological rhythms accordingly, which optimizes energy expenses (Bass et Takahashi, 2010; Green et al., 2008). The control of these oscillations involves transcriptional regulation by core clock genes encoding the transcription factors CLOCK, BMAL1, PER1, PER2, CRY1, and CRY2 (Takahashi, 2017). At the whole organism level, the clock system of the suprachiasmatic nucleus of the hypothalamus is entrained by the light/dark cycle via retinal innervation and acts as the master regulator. It synchronizes peripheral clocks through various neuronal, physiological, and hormonal signals (Bass et Takahashi, 2010; Green et al., 2008; Mohawk et al., 2012). However, tissues outside the suprachiasmatic nucleus contain local clocks and food intake acts as a synchronizing cue for peripheral clocks such as the liver clock (Koronowski et al, 2019) which is particularly sensitive to feeding (Damiola et al., 2000; Droin et al., 2021; Greenwell et al., 2019; Saini et al., 2013; Stokkan et al., 2001; Vollmers et al., 2009; Weger et al., 2021). Insulin production by pancreatic β-cells is under the control of a circadian clock and responsive to feeding (Perelis et al., 2016). Therefore, circulating insulin may contribute to the resetting of peripheral clocks upon nutritional challenges.

Epidemiological studies have linked circadian disruption with metabolic diseases in humans (Panda 2016; Perelis et al., 2016; Takahashi et al., 2008). Moreover, experimental studies have shown that experimental diabetes and diet-induced obesity reprogram the liver clock (Eckel-Mahan et al., 2013; Guan et al., 2018; Kohsaka et al., 2017; Panda 2016). In addition, experiments performed in cell culture indicate that insulin can synchronize clock gene expression in hepatocytes (Yamajuku et al., 2012). Moreover, insulin and IGF-1 regulate translation of PER clock proteins to entrain circadian rhythms with feeding *in vivo* (Crosby et al, 2019). Insulin and IGF-1 can signal through distinct pathways involving insulin receptor (IR) (Batista et al., 2019a). IR can directly translocate to the nucleus and form transcriptional complexes with RNA polymerase to control gene expression (Hancock et al., 2019). IR activation can also activate either AKT or Ras/MAP kinase signaling that ultimately influences the activity of several transcription factors involved in the control of hepatocyte growth and metabolism (Batista et al., 2019a) including activation of lipogenesis and inhibition of gluconeogenesis (Brown and Goldstein, 2008; Kubota et al., 2016; Li et al., 2010; Titchenell et al., 2017). Sterol regulatory element-binding transcription factor 1 (SREBP1c) is one such insulin-sensitive transcription factor and a potent regulator of lipogenic gene expression in hepatocytes downstream of IR signaling (Haas et al., 2012; Hegarty et al., 2005; Horton et al., 2002; Li et al., 2010). It has also been shown to interact with CRY1 to repress glucose production by promoting the degradation of forkhead box O1 (FOXO1) (Jang et al., 2016), a key transcription factor governing the expression of genes coding for gluconeogenic enzymes such as G6Pase (glucose-6-phosphatase) and PEPCK (phosphoenolpyruvate carboxykinase) (Dong et al., 2008; Matsumoto et al., 2007; O’Sullivan et al., 2015). In order to investigate the possible influence of IR on the liver clock *in vivo* we used a mouse model of conditional deletion of this receptor in hepatocytes, which have been shown to control liver chronophysiology in response to feeding (Guan et al., 2020) and are the main IR-expressing liver cell type (Michael et al., 2000).

## RESULTS AND DISCUSSION

### Hepatocyte-restricted deletion of IR induces time-sensitive defect in glucose homeostasis

First, we deleted IR specifically in hepatocytes in adult male mice (Figure 1A–C) carrying LoxP sites flanking the fourth exon of the IR gene and expressing Cre recombinase in the liver under the control of the transthyretin promoter (IR^hep−/−^) (Nemazanyy et al., 2015; Smati et al., 2020). In addition, inducible and hepatocyte-restricted deletion of IR led to severe glucose intolerance (Figure 1D,E) combined with hyperinsulinemia (Figure 1F) likely due to both increased insulin secretion and decreased insulin clearance, as observed in mice with constitutive deletion of the receptor (Michael et al., 2000). In addition, inducible and hepatocyte-restricted deletion of IR also led to decreased relative liver weight (Figure S1A) and reduced circulating HDL cholesterol (Figure S1B) which recapitulate some key phenotypic traits of mice constitutively lacking the insulin receptor in hepatocytes (Biddinger et al., 2008; Michael et al., 2000). Both insulinemia and glycemia were measured in the light phase (ZT8) and in the dark phase (ZT16) (ZT=Zeitgeber time), revealing increased hyperinsulinemia as well as hyperglycemia at night (Figure 1F,G) in animals fed *ad libitum*. At these two time points, we also measured contrasted effect of IR loss in hepatocytes on other circulating lipids and hepatic metabolites (Figure S2A,B). Moreover, we further confirmed that AKT phosphorylation in the liver is largely dependent on IR expression in hepatocytes (Haas et al., 2012) and peaks at the transition between night and day (ZT12) when feeding starts (Figure 1H,I).

**Figure 1:**
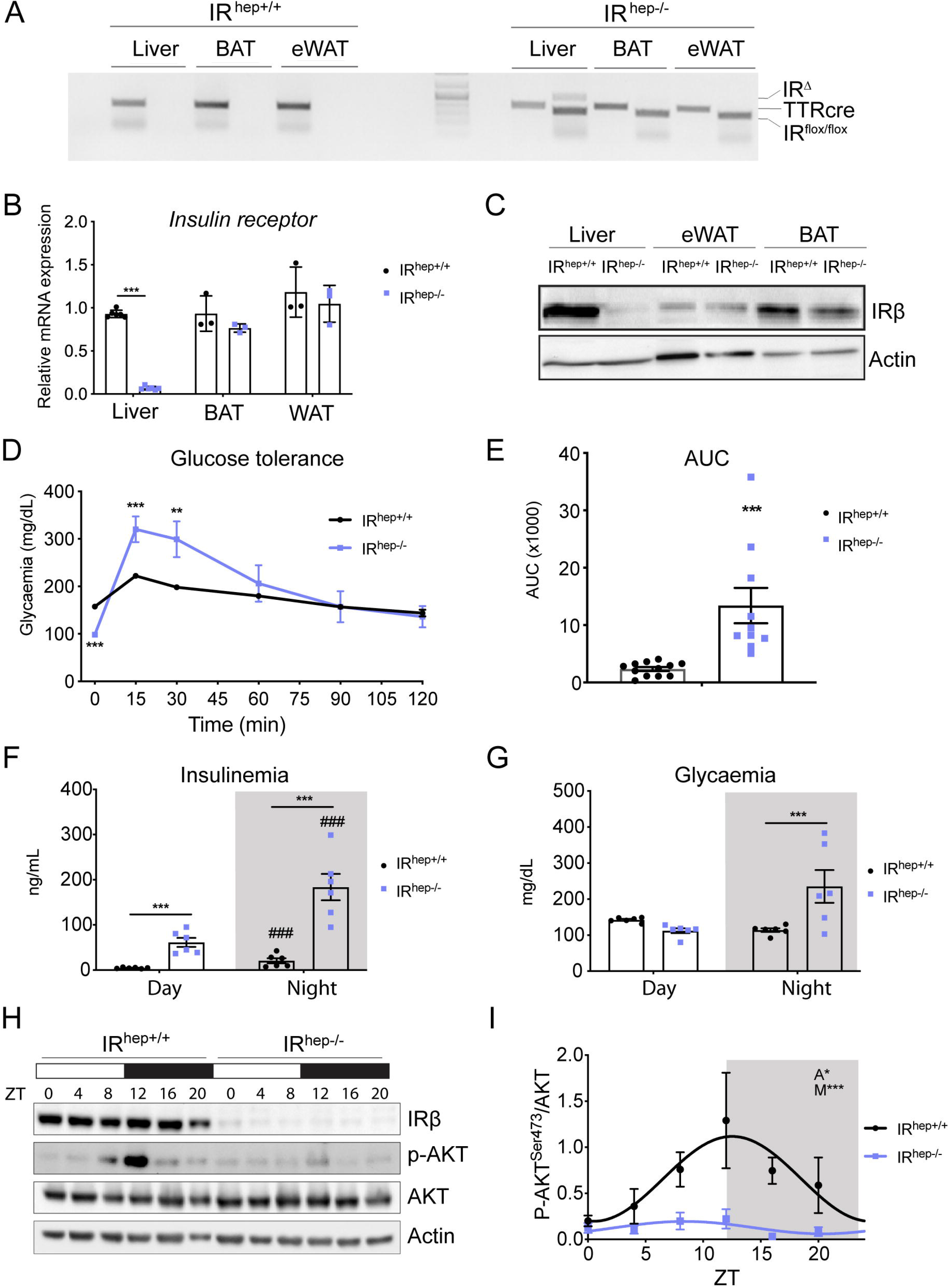
Hepatocyte-restricted deletion of the insulin receptor in adult mice leads to defective glucose homeostasis associated with impaired liver AKT signaling. **A**, PCR analysis of IR (insulin receptor) floxed (IR^hep+/+^) and TTR-^Tam^ -Cre (TTR-^Tam^-Cre^+/−^) genes from mice that are liver wild type (WT), (IR^hep+/+^) or liver knockout (IR^hep−/−^) for IR using DNA extracted from different organs (n=3 mice per group). **B**, Relative mRNA expression levels of IR from liver, epididymal white adipose tissue (eWAT), and brown adipose tissue (BAT) samples from IR^hep+/+^ and IR^hep−/−^ mice (n=3 mice per group). Data represent mean ± sem; statistical analysis was performed using two-way ANOVA with Sidak’s multiple comparison test. *p<0.05; **p<0.01; ***p<0.001 **C**, Insulin receptor-β was assessed by western blotting in liver, eWAT, and BAT (blots are representative of three independent experiments). **D,E**, Glucose (2g/kg) tolerance test after 6 h of fasting (**D**) and area under the curve (AUC) corresponding to the glucose tolerance test (**E**) (n=10 mice per group). Data represent mean ± sem; statistical significance was calculated using a two-tailed Student t-test with Bonferroni’s multiple comparisons. **F**, Quantification of circulating Insulin levels by ELISA (n=6 mice per group). Data represent mean ± sem; statistical analysis was performed using two-way ANOVA with Sidak’s multiple comparison test. # represents significant time effect and * represents significant genotype effect #,*p<0.05; ##,**p<0.01; ###,***p<0.001. **G**, Quantification of circulating glucose levels (n=6 mice per group). **H,I**, Insulin receptor-β, AKT phosphorylation, and total AKT were assessed by western blotting in liver (**H**) and relative quantification of AKT phosphorylation over total AKT (**I**) (blots are representative of five independent experiments. Data represent mean ± sem. Statistical analysis was performed using linear regression analysis fitting a 24 h cosinor model. A=amplitude; M=mesor.

### Hepatocyte-restricted deletion IR induces time-sensitive changes in liver gene expression

To assess the consequences of IR deletion on gene expression and its contribution of IR to the control of nocturnal and diurnal liver activity, we analyzed liver gene expression during the day (ZT8) and at night (ZT16). The expression of known insulin-sensitive genes such as *Gck* and *Pck1* was altered by IR deletion (Figure 2A and Figure S2C). Genome-wide expression profiles revealed a large set of genes (2389) which were differentially expressed between IR^hep+/+^ and IR^hep−/−^ mice at both ZT8 and ZT16 (Figure 2B and Figure S3D,E). These genes discriminated IR^hep+/+^ and IR^hep−/−^ mice when hierarchical clustering was performed (Figure 2C). They were grouped in three clusters (1, 5, and 6). Cluster 1 relates to inflammation that is elevated in response to IR deletion while clusters 5 and 6 are enriched in Srebp1c targets, which are metabolic genes downregulated in IR^hep−/−^ mice (Figure 2D and Figure S3). In liver of mice from both genotypes, 1773 genes were identified as differentially expressed between night and day (Figure 2B). Genes whose expression was not significantly impacted by the hepatocyte-restricted deletion of IR are included in clusters 4 and 8. Cluster 4 is enriched in genes associated with functions such as DNA repair, which is downregulated during the night (Figure 2E and Figure S3). Conversely, cluster 8 is enriched in genes up-regulated during the night which are associated with functions such as protein folding (Figure 2E and Figure S3).

**Figure 2:**
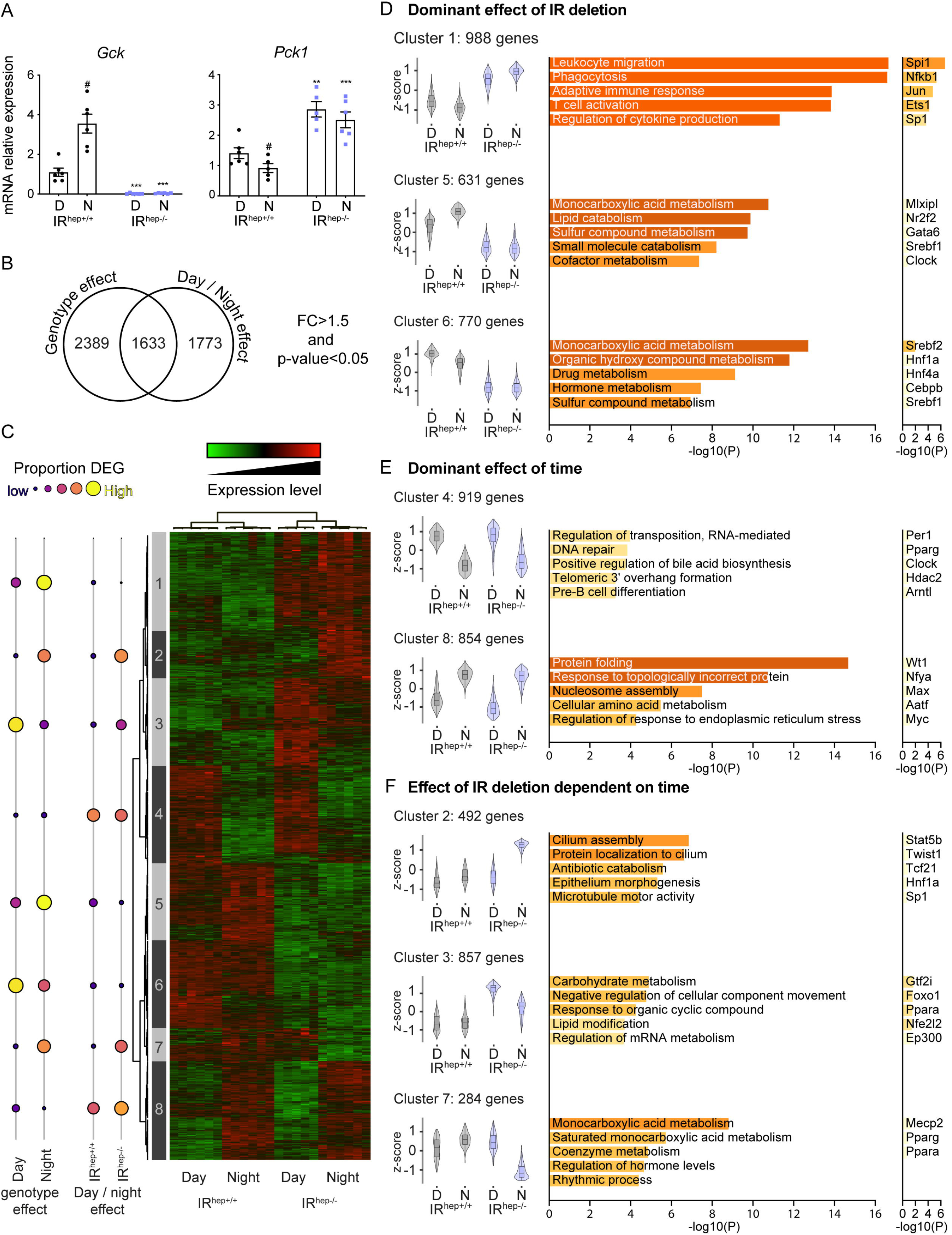
Hepatocyte-restricted deletion of the insulin receptor in adult mice induces time-sensitive changes in liver gene expression. Twelve-week-old IR^hep+/+^ and IR^hep−/−^ male mice were fed *ad libitum* and killed at ZT8 (day, D) or ZT16 (night, N). **A**, Relative mRNA expression levels of *Gck* and *Pck1* in liver samples (n=5–6 mice per group). Data represent mean ± sem; statistical analysis was performed using two-way ANOVA with Sidak’s multiple comparisons test. # represents significant time effect and * represents significant genotype effect #,*p<0.05; ##,**p<0.01; ###,***p<0.001. **B**, Venn diagram representing differentially expressed genes (DEG, p<0.05) influenced by genotype and time (Fold change >1,5). **C**, Heat map based on DEG: circles on the left represent the proportion of significantly DEG in each cluster, within each comparison (n=6 mice per group). **D-F**, Representative violin plot, gene ontology (GO) biological process, and enrichment of transcription factors (TTRUST) for each cluster(p-value cutoff<0.05; min enrichment 1,5), (**D**) dominant effect of IR deletion, (**E**) dominant effect of time, and (**F**) interaction between genotype and time.

Moreover, among the 3406 genes which were found differentially expressed between night and day, 1633 genes were differentially expressed between IR^hep+/+^ and IR^hep−/−^ mice (Figure 2B). These genes belong to clusters 2, 3, and 7 (Figure 2C-F and Figure S3). All three of these gene clusters have a similar pattern. In IR^hep+/+^ mice there were no obvious differences between their nocturnal and diurnal expression, but they became time-sensitive in IR^hep−/−^ mice (Figure 2F). Cluster 2 includes 492 genes related to the cilium structure and plasma membrane, with increased nocturnal expression in IR^hep−/−^ mice but not in IR^hep+/+^ mice. Cluster 3 includes 857 genes related to FOXO1 and other key signaling pathways involved in metabolic homeostasis. In agreement with the role of insulin signaling in the repression of FOXO-mediated regulation of gene expression, genes from this cluster were more expressed in IR^hep−/−^. Moreover, within this cluster, nocturnal expression is lower than diurnal expression in IR^hep−/−^ mice. Finally, cluster 7 includes 284 genes related to metabolism with expression reduced specifically during the night in IR^hep−/−^ mice. Altogether, these data establish that liver gene expression is both time-dependent and IR-sensitive. With the exception of cluster 1, IR-sensitive functions were mostly associated with metabolism. In hepatocytes, this IR-sensitive regulation of metabolic gene expression is generally considered to be regulated by PI3K-AKT signaling (Batista et al., 2019a). Downstream of AKT signaling, transcription factors such as SREBPs and FOXO1 are key players known to be involved in the metabolic response to insulin. Our results identify SREBP as belonging to a group of other influential transcription factors consistently controlling IR-sensitive gene expression both during the light phase and during the dark phase. By contrast, other transcription factors such as Foxo1 appear to be IR-sensitive only during the dark phase.

### Hepatocyte-restricted deletion of IR disrupts the rhythmicity of liver gene expression

To investigate the circadian control of IR-dependent metabolism, we analyzed blood insulin, glucose, and free fatty acid (FFA) concentrations every 4 hours around the clock in IR^hep+/+^ mice and IR^hep−/−^ mice fed *ad libitum* (Figure 3A and Figure S4A). Hepatocyte-restricted deletion of IR induced a time-sensitive increase in glycaemia, which was maximal at the end of the dark phase. Hyperinsulinemia was consistently observed in IR^hep−/−^ mice. FFA levels were not significantly impacted by hepatocyte IR deletion but tended to be higher in the light phase than during the dark phase. We next examined the hepatic expression of core clock genes as well as clock-controlled genes in IR^hep+/+^ and IR^hep−/−^ mice. Hepatocyte-restricted deletion of IR did not modify *Clock* expression but led to significant shifts in the rhythmic expression of some of the core clock genes such as *Per2, Cry1*, *Cry2 and Bmal1* as well as clock-controlled genes such as *Hlf*, *Pparα* and *Tef* (Figure 3B) and genes representative of liver functions such as *Hmgcs1* and *Fgf21* (Figure S4B). To further examine the potential consequences of such changes on circadian entrainment of the liver oscillation, we performed genome-wide analysis of the transcriptome in mice fed *ad libitum* after hepatocyte-specific deletion of IR. Our analysis identified three groups of genes with rhythmic expression. Cluster 1 was equally rhythmic in IR^hep+/+^ mice and IR^hep−/−^ mice (Figure 3C) and includes 646 genes associated with functions such as DNA repair that are enriched among the targets of the transcription factors Mecp2, E2f1, Clock, and AR (Figure 3F). Cluster 2 comprises 1393 genes showing disrupted oscillation in IR^hep−/−^ mice (Figure 3D). Their functions include chromosome segregation, autophagy of peroxisome and regulation of translation and are enriched in targets of Foxo1, Rbpj and Trp53 (Figure 3F). The expression of an additional cluster of 1295 genes (cluster 3) oscillated only in IR^hep−/−^ mice (Figure 3E). This cluster is enriched in genes with functions related to cell structure, metabolism, and division, and in targets of Sp1, Foxo4, Per1 (Figure 3F). Altogether, this analysis provides evidence that hepatocyte-restricted deletion of IR is sufficient to alter the circadian rhythmicity of gene expression in the liver. Importantly, IR deletion led not only to reduced rhythmicity but also induced gene expression reprogramming and novel oscillating transcripts.

**Figure 3:**
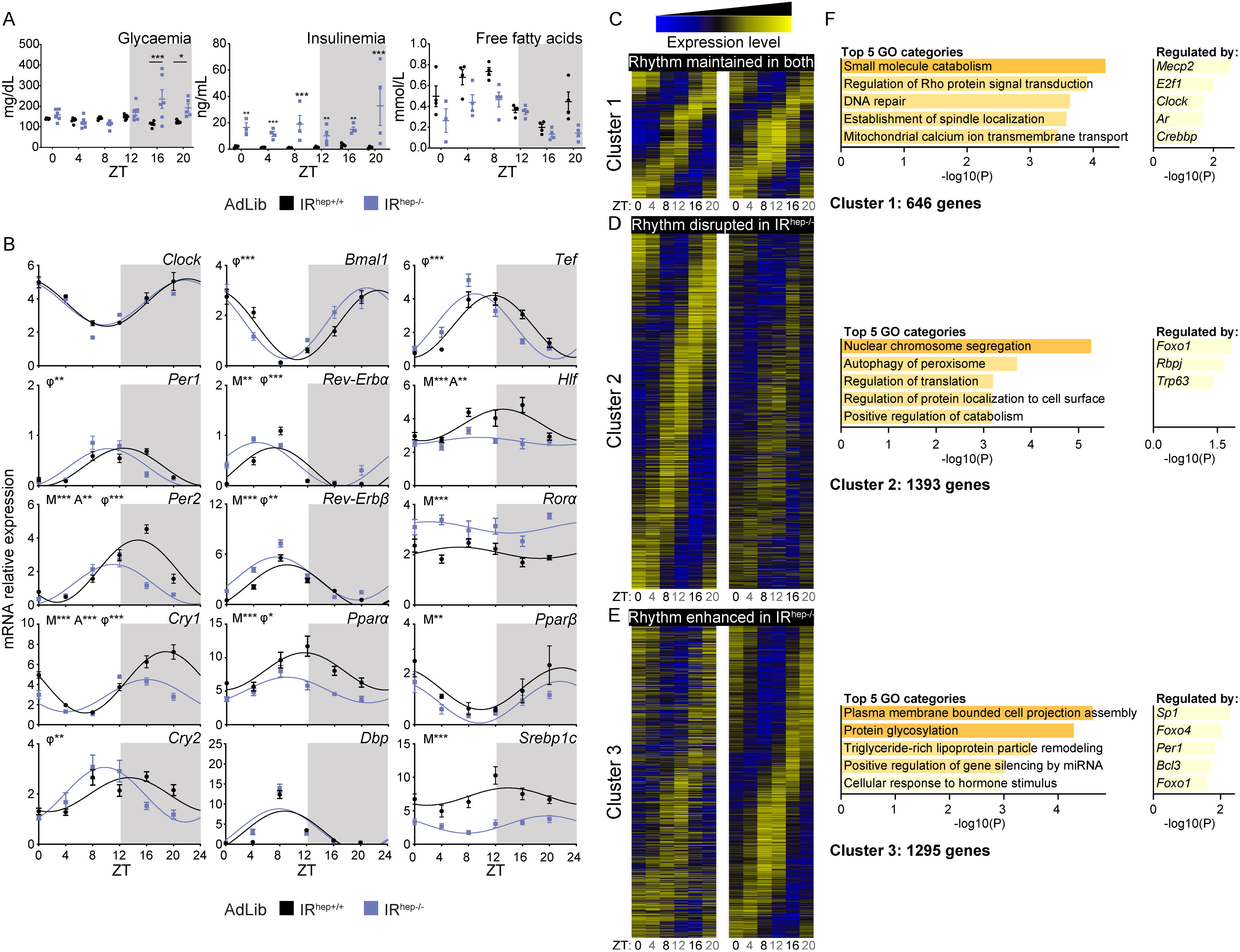
Hepatocyte-restricted deletion of the insulin receptor in adult mice disrupts the core clock and the rhythmicity of liver gene expression. Twelve-week-old IR^hep+/+^ and IR^hep−/−^ male mice were fed *ad libitum* and killed at different Zeitgeber times (ZT) from ZT0 to ZT20. **A**, Quantification of circulating glucose, insulin, and FFA levels (n=6 mice per group). Data represent mean ± sem; statistical significance was calculated using two-way ANOVA with Sidak’s multiple comparisons, *p<0.05; **p<0.01; ***p<0.001. **B**, Relative mRNA expression levels of core clock genes (*Clock*, *Per1*, *Per2*, *Cry1*, *Cry2*, *Bmal1*, *Rev-Erbα*, and *Rev-Erbβ*) and clock-controlled genes (*Pparα, Dbp*, *Tef*, *Hlf*, *Rorα, Pparβ* and *Srebp1c*) (n=6 mice per group). Data represent mean ± sem; statistical analysis was performed using linear regression fitting a 24-h cosinor model. A=amplitude; M=mesor; ϕ= accrophase, *p<0.05**; p<0.01; ***p<0.001. **C-E**, Heat maps of genes with rhythmic expression: (**C**) genes with rythmic expression in both genotypes, (**D**) genes with rhythmic expression disrupted only in *IR^hep−/−^*mice, and (**E**) genes with rhythmic expression enhanced only in *IR*^*hep*−/−^ mice (n=3 mice per group). **F**, Gene ontology (GO) biological process and enrichment of transcription factors (TTRUST) for each cluster (p-value cutoff <0.05; min enrichment 1,5).

### Hepatocyte IR is required for the reprogramming of liver gene expression rhythmicity in response to feeding

Food restriction during the night (ZT12–ZT24) has been shown to uncouple the liver clock from the central clock (Damiola et al., 2000). We used a food-restriction protocol to challenge the liver clock and further investigate the role of hepatocyte IR in its programming. Two weeks post induction of IR deletion, IR^hep+/+^ mice and IR^hep−/−^ mice were fed only during the light phase for 15 days (Figure 4A and Figure S4C). While hyperglycemia induced by IR deletion in mice fed *ad libitum* was maximal between ZT16 and ZT20, in mice subjected to restricted feeding it was maximal at ZT12 (Figure 4B). Unlike in *ad libitum* fed conditions, IR^hep−/−^ mice under food restriction did not display consistent hyperinsulinemia. Finally, FFA levels were higher during the light phase in *ad libitum* fed mice, with no significant difference between IR^hep+/+^ mice and IR^hep−/−^ mice. In contrast, in restricted-feeding (RF) mice, FFA levels were higher during the dark phase in mice of both genotypes, which is likely due to food restriction during the dark phase. Moreover, FFA levels in RF mice during the dark phase were significantly increased in IR^hep+/+^ mice when compared with IR^hep−/−^ mice.

**Figure 4:**
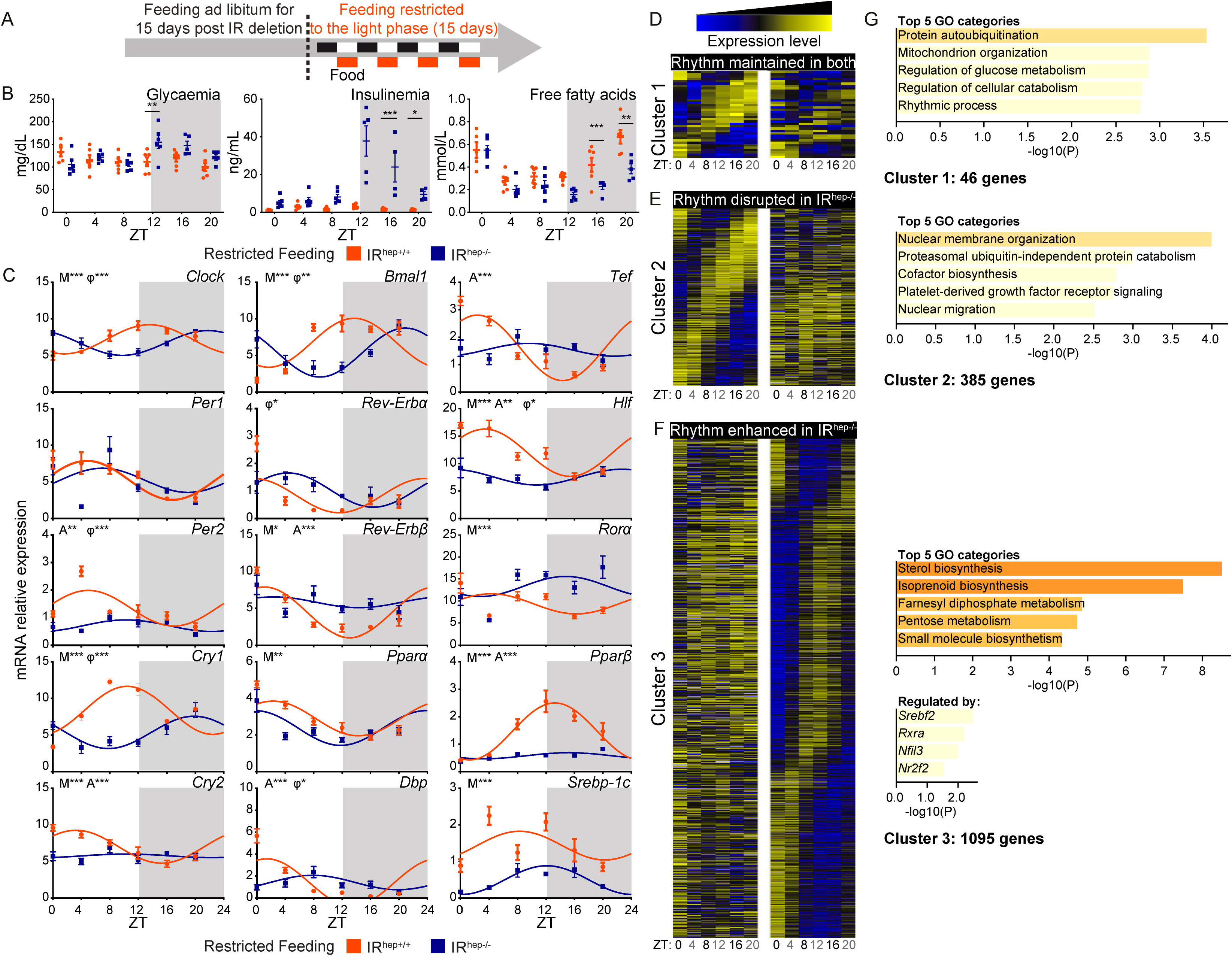
Hepatocyte-restricted deletion of the insulin receptor in adult mice disrupts the reprogramming of core clock and rhythmic liver gene expression in response to restricted feeding. **A**, Eight-week-old IR^hep+/+^ and IR^hep−/−^ male mice were fed ad libitum for 15 days after *IR* deletion, then fed during the light period only for 15 days (restricted feeding). **B**, Circulating glucose, insulin, and FFA levels were analyzed (n=6 mice per group). Data represent mean ± sem; statistical significance was calculated using two-way ANOVA with Sidak’s multiple comparisons, *p<0.05; **p<0.01; ***p<0.001. **C**, Relative mRNA expression levels of core clock genes (*Clock*, *Per1*, *Per2*, *Cry1*, *Cry2*, *Bmal1*, *Rev-Erbα* and *Rev-Erbβ*) and clock-controlled genes (*Pparα, Dbp*, *Tef*, *Hlf*, *Rorα*, *Pparβ* and *Srebp1c*) (n=6 mice per group). Data represent mean ± sem; statistical analysis was performed using linear regression analysis fitting a 24-h cosinor model. A=amplitude; M=mesor; ϕ=accrophase, *p<0.05, **p<0.01, ***p<0.001. **D-G**, Heat maps of rhythmically expressed genes and associated transcription factors and GO enrichment: (**D**) genes with rhythmic expression in both genotypes, (**E**) rhythmic genes disrupted in *IR^hep−/−^* mice, and (**F**) rhythmic genes enhanced only in *IR^hep−/−^* mice with (n=3 mice per group). **F**, Gene ontology (GO) biological process and enrichment of transcription factors (TTRUST) for each cluster (p-value cutoff <0.05; min enrichment 1,5).

The expression of core clock genes and clock-controlled genes was modified under restricted feeding and peaked with a 10–12 hour shift when compared with their *ad libitum* peaks (Figure 4C). Surprisingly, mice with hepatocyte-restricted IR deletion were refractory to RF-mediated adaptation of the core clock gene expression in the liver. To further analyze the consequences of such defective modification of the clock, we analyzed the rhythmicity of gene expression under restricted feeding. We identified 3 groups of genes that were rhythmic in mice of both genotypes, or in IR^hep+/+^ or IR^hep−/−^ mice. Only 46 genes were present in both groups (Figure 4D). A larger cluster of 385 genes was rhythmic only in IR^hep+/+^ mice (Figure 4E). While these two clusters relate to specific gene annotations, they were not significantly enriched in transcription factor targets. Finally, a large cluster of 1095 genes was rhythmic only in the liver of mice lacking IR (Figure 4F). This was confirmed by measuring the expression of a set of hepatic genes such as *Igfbp1*, *Fgf21*, *Pcsk9* and *G6pc* (Figure S4D). In that cluster, functions associated with glucose and lipid metabolism were overrepresented, consistent with the enrichment in targets of Srebp2, Rxrα, and Nfil3. Under restricted feeding, novel oscillations in liver gene expression occurred with strong dependence on the expression of IR in hepatocytes. Overall, under restricted feeding, rhythmic reprogramming was reduced and the rhythmicity of novel oscillations was enhanced in mice lacking hepatocyte IR.

## Conclusions

Our work extends previous findings regarding the role of IR in the control of liver gene expression (Batista et al., 2019a; Batista et al., 2019b; Haas et al., 2012; Titchenell et al., 2017) and provides *in vivo* evidence that IR-dependent signaling in hepatocytes synchronizes the liver clock. The loss of IR in hepatocytes led to both the loss of pre-existing oscillations and a newly enhanced rhythmicity, which are typically observed after genetic manipulation of the core clock genes in the liver and in diet-induced reprogramming of liver gene expression (Eckel-Mahan et al., 2013; Guan et al., 2018; Kohsaka et al., 2017; Panda et al., 2016). There are three mechanisms by which IR influences gene expression in response to insulin (and insulin-like growth factors): direct translocation to the nucleus and activation of either AKT-dependent or Ras/MAP kinase-dependent signaling (Batista et al., 2019a; Batista et al., 2019b). Further research is required to define which of them is responsible for programming the liver clock and whether it also regulates post-transcriptional (Sinturel et al., 2017; Wang et al., 2018), translational (Atger et al., 2015; Jouffe et al., 2013; Mauvoisin et al., 2014; Sinturel et al., 2017) or post-translational events (Mauvoisin et al., 2014; Mauvoisin et al., 2017; Mauvoisin et Gachon, 2020).

It is well known that feeding acts as a predominant signal within the liver to reprogram the clock and the rhythmicity of gene expression (Damiola et al., 2000; Greenwell et al., 2019; Saini et al., 2013; Stokkan et al., 2001; Vollmers et al., 2009). Our work sheds light on the physiological importance of intact hepatocyte IR for the oscillation of gene expression in response to feeding. In the liver, several types of nutritional challenge can lead to significant resetting of the clock. Diet-induced obesity (Eckel-Mahan et al., 2013; Guan et al., 2018; Kohsaka et al., 2017; Quagliarini et al., 2019) and time-restricted feeding (Saini et al., 2013; Stokkan et al., 2001; Vollmers et al., 2009) both influence the liver clock. Our findings support the hypothesis that such metabolic challenges that affect insulin signaling may, at least in part, reprogram the liver clock through changes in hepatocyte insulin sensitivity.

In addition, our results support a role for liver insulin signaling in clock control that may be shared with other insulin-sensitive tissues. Therefore, defective insulin signaling may contribute directly to the disruption of circadian homeostasis that occurs in diabetes and its complications.

## STAR METHODS

### KEY RESSOURCES TABLE

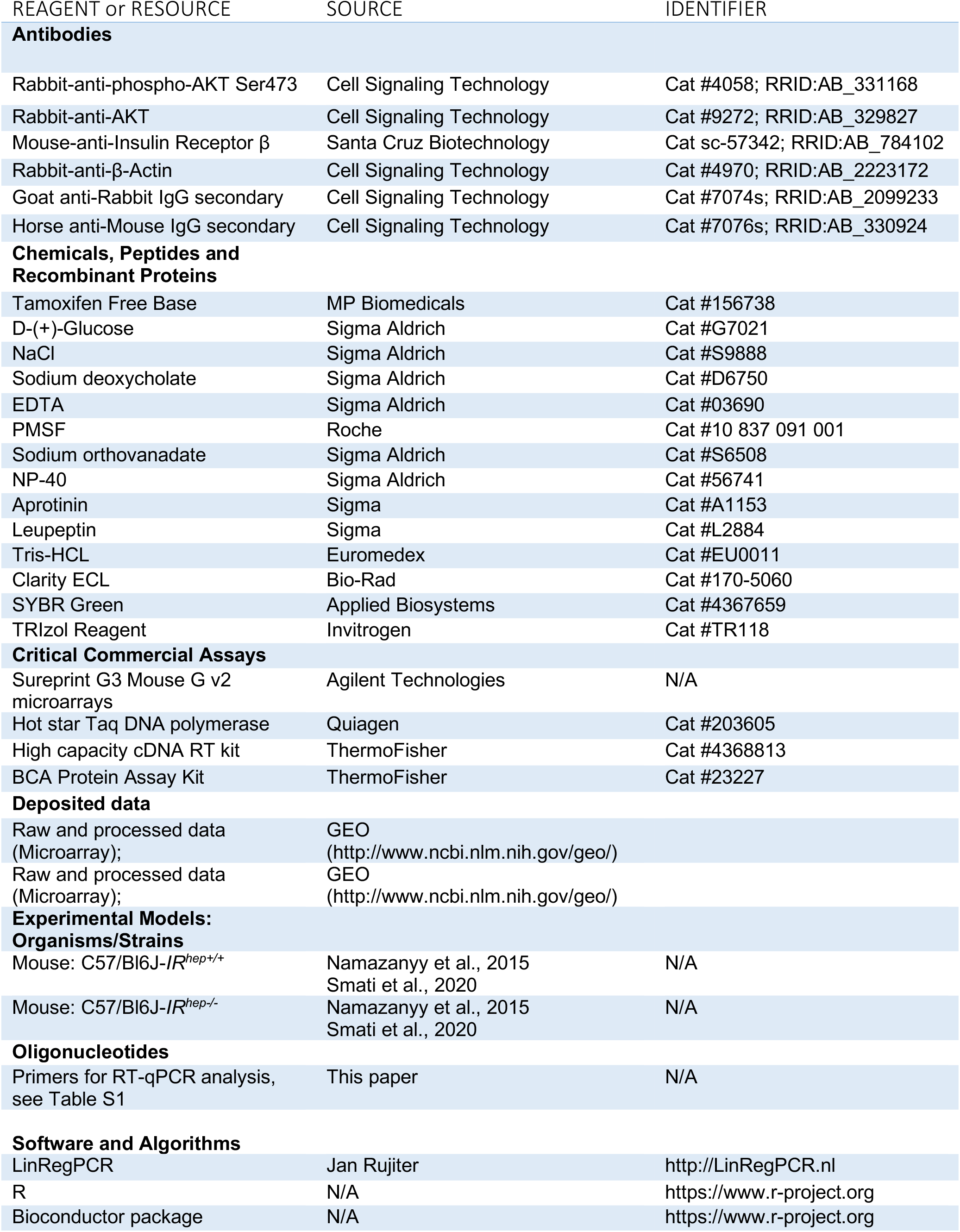
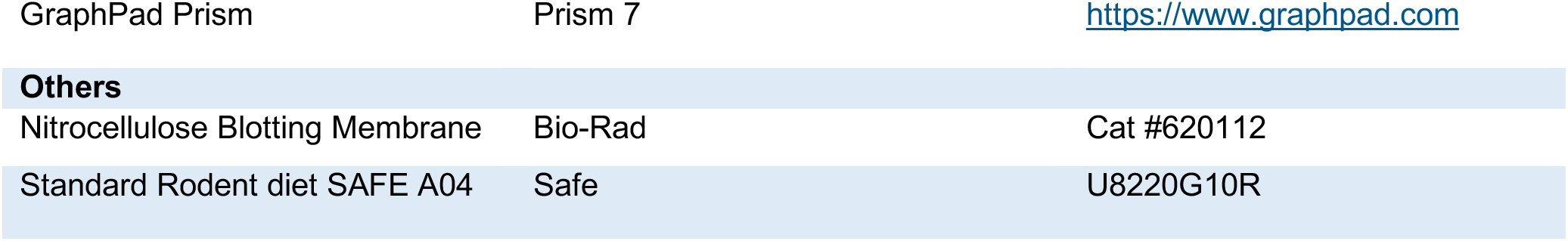

### CONTACT FOR REAGENT AND RESOURCE SHARING

Further information and requests for resources and reagents should be directed to and fulfilled by the lead contact, Hervé Guillou.

## EXPERIMENTAL MODEL

### *In vivo* studies

In vivo studies were performed in accordance with European guidelines for the use and care of laboratory animals, and approved by an independent Ethics Committee. To generate tamoxifen-inducible hepatocyte-specific IR knockout mouse line (IR^hep−/−^) and their control (IR^hep+/+^), animals carrying LoxP sites flanking the fourth exon of the IR gene (IRlox/lox stock number: 006955; Jackson Laboratory, Bar Harbor, ME, USA) were intercrossed with C57BL/6J mice, which specifically express Cre recombinase in the liver under the transthyretin promoter (TTR-CreTam mice), as previously described^32,33^.

At the age of 8 weeks, males IR^hep−/−^ (IR^floxed/floxed / TTR-CreTam+/+)^) and IR^hep+/+^ (IR^floxed/floxed / TTR-CreTam−/−^) mice received tamoxifen (Tamoxifen Free Base, MP Biomedicals) by intraperitoneal injection (1,5mg/kg) during three consecutive days, in order to induce deletion of hepatocyte Insulin receptor. First experiments were started two weeks after tamoxifen injection.

The IR deletion was confirmed in IR^hep−/−^ mice with PCR and HotStar Taq DNA Polymerase (5 U/μl, Qiagen) using the following primers: IR-forward: 5’-GATGTGCACCCCATGTCTG-3’; IR-reverse: 5’-CTGAATAGCTGAGACCACAG-3’ and IRdelta: 5’-GGGTAGGAAACAGGATGG-3’. The amplification conditions were as follows: 95°C for 5 min, followed by 35 cycles of 94°C for 30 sec, 65°C for 30 sec, and 72°C for 45 sec, and a final cycle of 72°C for 7 min. This reaction produced 300-bp, which represented the IR sequence with the floxed allele. The TTR-CreTam allele was detected by PCR using the following primers: forward: 5′- CCTGGAAAATGCTTCTGTCCG-3′ and reverse: 5′- CAGGGTGTTATAAGCAATCCC-3′.

All mice were fed a standard rodent diet (Safe 04 U8220G10R), and housed under controlled temperature (21-23°C) and light (12-h light/12-h dark) conditions : lights were turned on at Zeitgeber time (ZT) 0 and turned off at ZT12.

For restricted feeding, mice were adapted during 2 weeks to eat during the light period, and food was removed at the beginning of the dark period, as previously described^12^.

### METHOD DETAILS

#### Blood and tissue sampling in mice

Prior to sacrifice, blood was collected into EDTA coated tubes (BD Microtainer, K2E tubes) from the submandibular vein. All mice were killed in a fed state (unless stated otherwise). Plasma was collected by centrifugation (1500 xg, 15min, 4°C) and stored at −80°C. Following killing by cervical dissociation, organs were removed, weighted, dissected and used for snap frozen in liquid nitrogen and stored at −80°C.

#### Oral glucose tolerance test

OGTT was performed one month after tamoxifen injection. Mice were fasted for 6 h and received an oral (2g/kg body weight) glucose load (G7021, Sigma Aldrich). Blood glucose was measured at the tail vein using an AccuCheck Performa glucometer (Roche Diagnostics) at −15, 0, 15, 30, 45, 60, 90, and 120 minutes.

#### Immunoblotting

Frozen liver, white and brown adipose tissue samples, were homogenized in a lysis buffer [50 mM Tris-HCl (pH 7,4), 150 mM sodium chloride (NaCl), 1% NP-40, 0,25% sodium deoxycholate, 0,1% sodium dodecyl sulfate (SDS), 2 mM ethylenediaminetetraacetic acid (EDTA) and 1 mM phenylmethylsulfonyl fluoride (PMSF)] supplemented with 2 μg/mL aprotinin, 5 μg/mL leupeptin and 1 mM sodium orthovanadate for 30 min at 4°C. Samples were then centrifuged at 13,000 rpm for 30 min at 4°C. Supernatants were collected and the protein concentration in each lysate measured using BC Assay Protein Quantification Kits (Interchim, Montluçon, France). Proteins (20-30 μg) were separated by SDS-PAGE (polyacrylamide gel electrophoresis) and transferred onto nitrocellulose membranes, which were first incubated overnight at 4°C with primary antibodies against Insulin receptor β (sc-57342, Santa Cruz Biotechnology, Santa Cruz, CA, USA), pSer473AKT (#4058, Cell Signaling Technology, Beverly, MA, USA), total AKT (#9272, Cell Signaling Technology, Beverly, MA, USA) and β-actin (#4970, Cell Signaling Technology, Beverly, MA, USA), and then with horseradish peroxidase-conjugated secondary antibodies (anti-mouse for Insulin receptor β detection and anti-rabbit for all others) for 1 h at room temperature. Immunoreactive proteins were detected with Clarity Western ECL blotting substrate (Bio-Rad Laboratories, Hercules, CA, USA) according to the manufacturer’s instructions, and signals were acquired using the Chemidoc Touch Imaging System (Bio-Rad).

#### Plasma analysis

Plasma samples were assessed for free fatty acids, triglycerides, total cholesterol and low-density lipoprotein (LDL) cholesterol levels, using a Cobas Mira Plus biochemical analyzer (Roche Diagnostics, Indianapolis, IN, USA), at the ANEXPLO/ Genotoul Zootechnics Core Facilities in Toulouse. Plasma Insulin and Glucagon concentration was assessed using Insulin mouse serum Assay HTRF Kit (Cisbio) and Glucagon mouse serum Assay HTRF kit (Cisbio), at the WE-MET in Toulouse. Blood glucose was measured with an Accu-Chek Go glucometer (Roche Diagnostics).

#### ^1^H-NMR based metabolomics

Liver polar extracts were prepared for NMR analysis as described previously (Regnier et al., 2020). All ^1^H-NMR spectra were obtained on a Bruker DRX-600-Avance NMR spectrometer (Bruker, Wissembourg, France) using the AXIOM metabolomics platform (MetaToul) operating at 600.13 MHz for 1H resonance frequency using an inverse detection 5-mm 1H-13C-15N cryoprobe attached to a cryoplatform (the preamplifier cooling unit).

#### RNA extraction and RT-qPCR

Total cellular RNA was extracted using TRI Reagent (Molecular Research Center, Inc., Cincinnati, OH, USA). Total RNA samples (2 μg) were reverse-transcribed using High Capacity cDNA Reverse Transcription Kits (Thermo Fisher Scientific, Waltham, MA, USA) for real-time quantitative polymerase chain reaction (qPCR) analyses; the primers used for the SYBR Green assays are listed in Extended data Table 1. Amplifications were performed using a Stratagene Mx3005P system (Agilent Technologies, Santa Clara, CA, USA). The qPCR data were normalized to the level of *Psmb6* (proteasome 20S subunit beta 6) transcripts, and analyzed using LinRegPCR.v2015.3 to get the starting concentration (N0) which is calculated as follow: N0 = threshold/(Eff meanCq) with Eff mean: mean PCR efficiency and Cq: quantification cycle.

#### Microarray

Gene expression profiles were obtained at the GeT-TRiX facility (GenoToul, Genopole Toulouse Midi-Pyrénées) using Agilent Sureprint G3 Mouse GE v2 microarrays (8×60K, design 074809) following the manufacturer’s instructions.

For each sample, Cyanine-3 (Cy3) labelled cRNA was prepared from 200 ng of total RNA using the One-Color Quick Amp Labeling kit (Agilent) according to the manufacturer’s instructions, followed by Agencourt RNAClean XP (Agencourt Bioscience Corporation, Beverly, Massachusetts). Dye incorporation and cRNA yield were checked using a Dropsense™ 96 UV/VIS droplet reader (Trinean, Belgium). A total of 600 ng of Cy3-labelled cRNA was hybridized on the microarray slides following the manufacturer’s instructions. Immediately after washing, the slides were scanned on an Agilent G2505C Microarray Scanner using Agilent Scan Control A.8.5.1 software and the fluorescence signal extracted using Agilent Feature Extraction software v10.10.1.1 with default parameters. Microarray data and experimental details are available in NCBI’s Gene Expression Omnibus (Edgar, Domrachev, & Lash, 2002) and are accessible through GEO Series accession numbers GSE165154 and GSE165155.

### STATISTICAL ANALYSIS

#### Microarray data analyses

Microarray data were analyzed using R (R Development Core Team, 2008) and Bioconductor packages: www.bioconductor.org, v 3.0, (Gentleman et al., 2004) as described in GEO accession GSE165156. Raw data (median signal intensity) were filtered, log2 transformed, corrected for batch effects (microarray washing bath and labeling serials) and normalized using quantile method (Bolstad, et al. 2003).

A model was fitted using the limma lmFit function (Smyth, 2004). Pair-wise comparisons between biological conditions were applied using specific contrasts to extract genotype, Time and Genotype:Time interaction effects. A correction for multiple testing was applied using Benjamini-Hochberg procedure (BH, Benjamini et al. 1995) to control the False Discovery Rate (FDR). Probes with FDR ≤ 0.05 were considered to be differentially expressed between conditions.

Hierarchical clustering was applied to the samples and the differentially expressed probes using 1-Pearson correlation coefficient as distance and Ward’s criterion for agglomeration. The clustering results are illustrated as a heatmap of expression signals.

Rhythmicity analysis was performed with three R packages: MetaCycle (Wu et al., 2019), HarmonicRegression (Lück et al., 2015) and RAIN (Thaben et al., 2014). Program-specific settings were as follows: MetaCycle (meta2d): minper = 8, maxper = 24, cycMethod = “JTK”, adjustPhase = ‘predictedPer’, combinePvalue = ‘fisher’ ; HarmonicRegression: normalize = FALSE ; RAIN: period = 24, deltat = 4, nr.series = 3, peak.border = c(0.2, 0.8), method = ‘independent’. Resulting p values were adjusted using the Benjamini-Hochberg method within the p.adjust function available in base R to control for the false-discovery rate (FDR). Genes that were found to be rhythmic (q-value <0.01 and oscillation period within the range of 21 to 24 hr) in the 3 rhythmic programs were considered as rhythmic for that specific experimental condition.

#### Other statistical analyses

Statistical analyses were performed using GraphPad Prism version 8 for Mac OS X (GraphPad Software, San Diego, CA). Two-way ANOVA was performed followed by appropriate post-hoc tests (Sidak’s multiple comparisons test). P< 0.05 was considered significant.

Multiple T-test was also performed followed by appropriate correction (Benjamini, Krieger and Yekutieli). P< 0,05 was considered significant.

For circadian analysis, Nonlinear Regression (sine wave with nonzero baseline) was performed followed by extra sum-of-squares F test in order to compare amplitude (A), PhaseShift (φ) and Baseline (M= Midline estimating statistic of rhthym). Differences were considered statistically significant when P< 0,05.

### Data availability

All the data are available on request and gene expression analysis are available on GEO (GSE165154; GSE165155; GSE165156).

## Acknowledgements

We thank all members of the EZOP staff for their help during this project. We thank the staff from Genotoul (Anexplo, GeT-TRiX, Metatoul-Axiom) facilities for their help during this project.

