## Supplementary figures and images for "The hepatocyte insulin receptor is required to program rhythmic gene expression and the liver clock"

### Supplementary figure 1

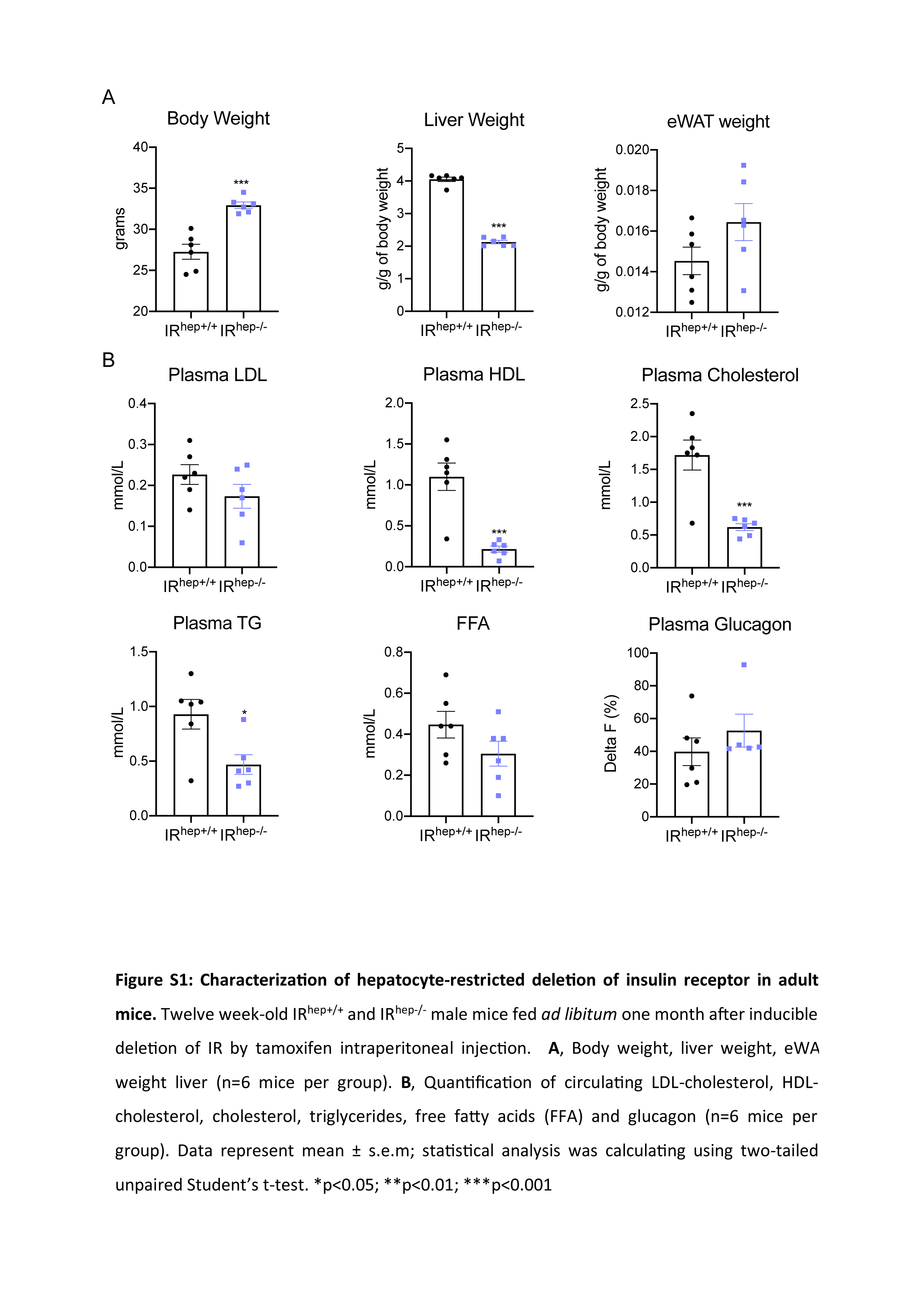

### Supplementary figure 2

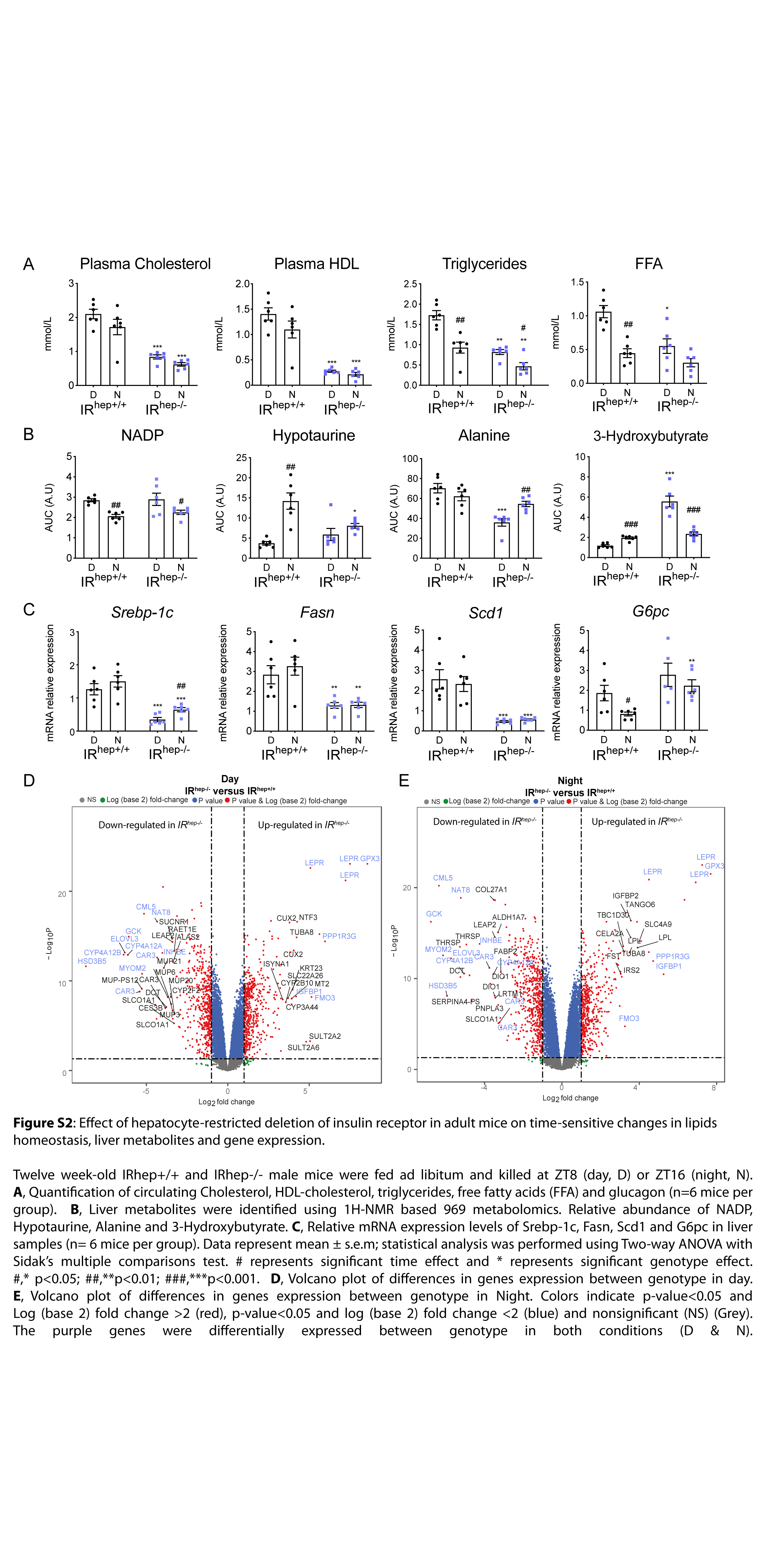

### Supplementary figure 3

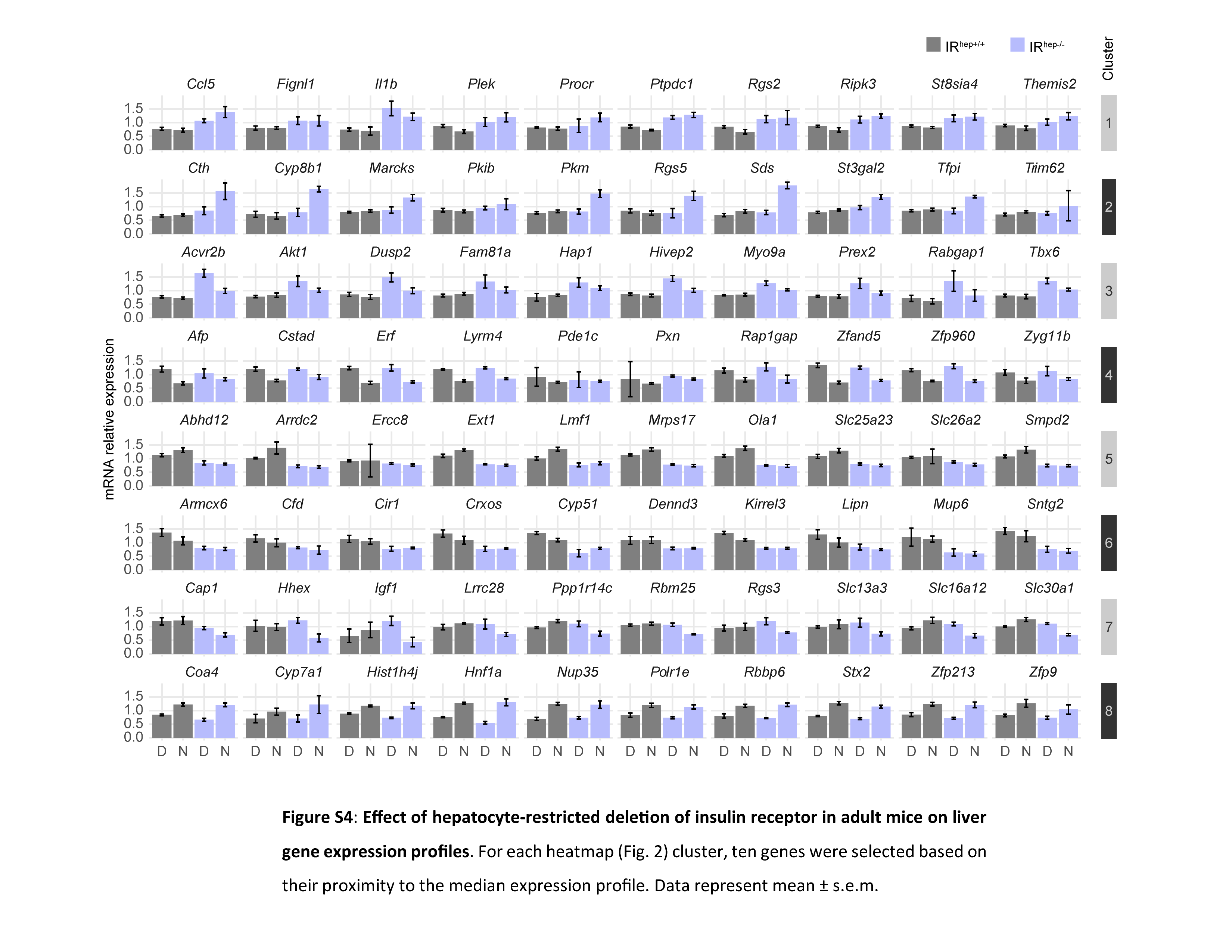

### Supplementray figure 4

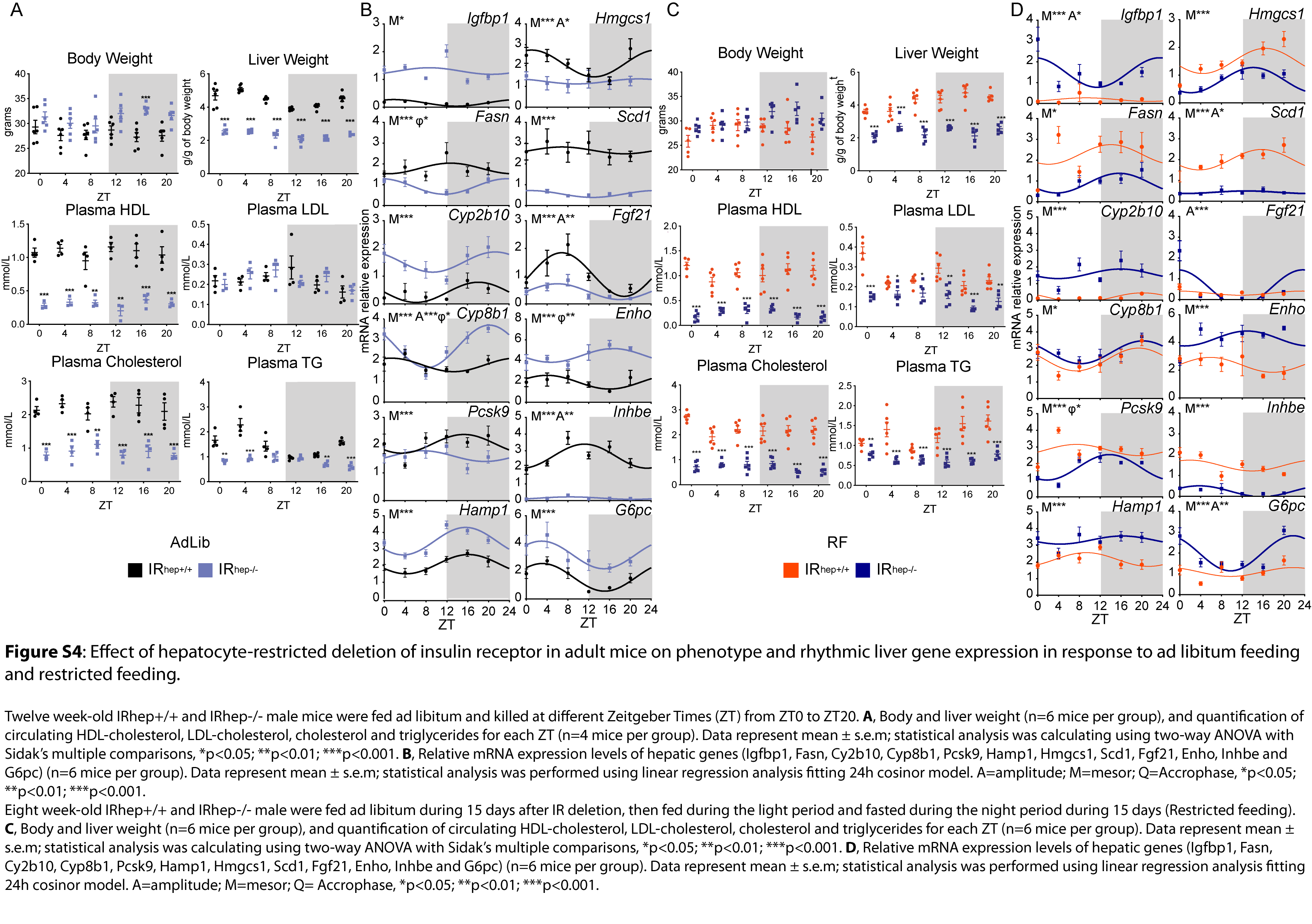
