## Supplementary table 1 for "The hepatocyte insulin receptor is required to program rhythmic gene expression and the liver clock"

***Extended data Table 1 : List of primers***

| Gene | NCBI Refseq | Forward primer | Reverse primer |
|---|---|---|---|
| *Bmal1* | NM_007489 | CAAACTACAAGCCAACATTTCTATCAG | TCGGTCACATCCTACGACAAAC |
| *Clock* | NM_007715 | GCTACAGTCACTACGTTCACTCAGG | CAAGCTACAGGAGCAGTCACTAATTT |
| *Cry1* | NM_001374058.1 | CTGGCGTGGAAGTCATCGT | CTGTCCGCCATTGAGTTCT |
| *Cry2* | NM_009963.4 | ACCGATGGAGGTTCCTACTG | AGCCTTGGGAACACATCAG |
| *Cyp2b10* | NM_009999 | TTTCTGCCCTTCTCAACAGGAA | ATGGACGTGAAGAAAAGGAACAAC |
| *Cyp8b1* | NM_010012 | CTCAAGGTGTTTGGGTACCAGTC | TCAGATGCTTGGTACTAGCTAAGTGG |
| *Dbp* | NM_016974.3 | AAGAAGGCAAGGAAAGTCCA | TGTACCTCCGGCTCCAGTA |
| *Enho* | NM_027147 | ACCGGGCTCAACTCAGGC | TGGCTGTCCTGTCCACACAC |
| *Fasn* | NM_007988 | AGTCAGCTATGAAGCAATTGTGGA | CACCCAGACGCCAGTGTTC |
| *Fgf21* | NM_020013.4 | AAAGCCTCTAGGTTTCTTTGCCA | CCTCAGGATCAAAGTGAGGCG |
| *G6pc* | NM_008061 | CTCACTTTCCCCACCAGGTC | GCTGAAAGTTTCAGCCACAGC |
| *GCK* | NM_010292 | TCGCAGGTGGAGAGGGA | TCGCAGTCGGCGACAGA |
| *Hamp1* | NM_032541.2 | AAGCAGGGCAGACATTGCG | CAGGATGTGGCTCTAGGCT |
| *Hlf* | NM_172563.3 | CGTCTCCGAACTGTATGCAGAG | GGTCAATGGGACTCGGTGTATT |
| *Hmgcs1* | NM_145942.4 | TCAATGCCGTAACTGGGT | CAATGTCTCCTGCAACTACCAGAG |
| *Igfbp1* | NM_008341.4 | CCTGCCAACGAGAACTCTAT | AGGATTTTCTTTCCACTCC |
| *Inhbe* | NM_008382.3 | TCAGCTTTGCTACCATCATAGACA | CATGGAGCGGTAGGTTGAAGT |
| *Insr* | NM_010568.2 | CACTGTCATCAATGGGCAGTTT | CATCAGGTTCCGAACAGTTGC |
| *Pck1* | NM_011044 | GAACCCCAGCCTGCCC | GAGCAACTCCAAAAAACCCG |
| *Pcsk9* | NM_153565 | AGGAAGACCGCTCCCCTG | TGGTATCTAAGAGATACACCTCCACCT |
| *Per1* | NM_011065.5 | ACCAGCGTGTCATGATGAC | CTCTCCCGGTCTTGCTTCAG |
| *Per2* | NM_011066.3 | ATGCTCGCCATCCACAAGA | GCGGAATCGAATGGGAGAA |
| *Ppar*$\alpha$ | NM_011144 | CCCTGTTTGTGGCTGCTATAATTT | GGGAAGAGGAAGGTGTCATCTG |
| *Ppar*$\beta$ | NM_011145 | AAGTGGCCATGGGTGACG | TGGTCCAGCAGGGAGGAAG |
| *Rev-Erb*$\alpha$ | NM_145434 | CAGCTGGTGAAGACATGACGAC | GGAGGAGCCACTAGAGCCAA |
| *Rev-Erbβ* | NM_011584 | CGCCATGGAGCTGAACG | GACAAGAGGCAGGGCTGGA |
| *Scd1* | NM_009127 | CAGTGCCGCGCATCTCTAT | CAGCGGTACTCACTGGCAGA |
| *Srebp-1c* | NM_01148 | GGAGCCATGGATTGCACATT | GCTTCCAGAGAGGAGGCCAG |
| *Tef* | NM_017367.3 | GCCGAGCTTCGCAAGGA | ACAGGTTACAAGGGCCCGT |
